# LineageSim: A Single-Cell Lineage Simulator with Fate-Aware Gene Expression

**DOI:** 10.64898/2026.02.08.704635

**Authors:** Haizhi (Gary) Lai, Mehrshad Sadria

## Abstract

Single-cell lineage data paired with gene expression are critical for developing computational methods in developmental biology. Since experimental lineage tracing is often technically limited, robust simulations are necessary to provide the ground truth for rigorous validation. However, existing simulators generate largely Markovian gene expression, failing to encode the fate bias observed in real biological systems, where progenitor states exhibit early signatures of future commitment. Consequently, they cannot support the training and evaluation of computational methods that model long-range temporal dependencies. We present LineageSim, a generative framework that introduces fate-aware gene expression, where progenitor states carry latent signals of their descendants’ terminal fates. This framework establishes a new class of benchmarks for cell fate prediction algorithms. We validate the presence of these temporal signals by training a logistic regression baseline, which achieves 68.3% balanced accuracy. This confirms that the generated data contain subtle but recoverable fate information, in contrast to existing simulators, where such predictive signals are systematically absent.

## 1 Introduction

Developmental biology aims to understand how progenitor cells differentiate into specialized cell types. An ideal experiment would map the complete cell lineage tree and track each cell’s gene expression over time. While state-of-the-art methods combine lineage tracing (heritable barcodes) with single-cell RNA sequencing (scRNA-seq), “ground truth” data remain elusive. Experimental limitations, such as the sparsity of lineage scarring and the destructive nature of sequencing, prevent us from observing the complete lineage history and the longitudinal gene expression of a single cell. Barcoding technologies often suffer from stochastic silencing or dropout, leaving large gaps in the reconstructed phylogenies. Sequencing requires lysing the cell, forcing us to infer dynamic temporal trajectories from static snapshots [Kester and van Oudenaarden, 2018]. As a result, real-world datasets are fragmented snapshots with noisy, incomplete labels, making the validation of computational methods notoriously difficult.

To address these gaps, simulation tools like TedSim [Pan et al., 2022] generate synthetic ground truth for benchmarking trajectory inference. However, these tools fail to capture a critical biological property: fate bias, the early transcriptional signatures in progenitor cells that predict future lineage, detectable before cell division [Wang et al., 2022, Sadria and Bury, 2024, Sadria and Swaroop, 2025]. Modeling fate bias requires long-range dependencies, as progenitor expression must carry information about distant descendants. Existing simulators lack this capability, generating expression primarily based on immediate state. Current benchmarks are therefore unsuitable for evaluating fate prediction methods: the synthetic data lack the very signal these models aim to learn.

We introduce LineageSim, a single-cell lineage simulator that generates fate-aware gene expression. By injecting a fate bias into the latent representations that determine expression, LineageSim generates lineage data where each progenitor’s expression encodes information about its descendants’ terminal fates. This approach captures the fate bias observed in real biological systems, enabling benchmarking of cell fate prediction methods [Sadria et al., 2024], which was not possible with previous simulators. We validate that these fate signals are detectable: a logistic regression classifier trained on early progenitor expression predicts terminal fate at 68.3% balanced accuracy, well above chance level. LineageSim is released as an open-source Python package designed for reproducibility and ease of use.

## 2 Method

LineageSim generates a lineage tree of cells and their gene expression in four phases.

### Phase 1: Tree generation

We generate the phylogenetic tree structure via stochastic cell division simulation.

### Phase 2: Fate assignment

After tree generation, we designate a commitment depth *D* at which fate decisions occur and probabilistically assign a binary cell fate (0 or 1) to each cell at that depth. The class probability is configurable, defaulting to 50/50. Each cell’s assigned fate is propagated to all of its descendants. Crucially, we then calculate a fate bias *p* for each progenitor, defined as the fraction of its committed descendants belonging to Fate 1, yielding a continuous value *p* ∈ [0, 1] that encodes its future developmental trajectory.

### Phase 3: Latent Manifold Generation

We simulate a latent representation vector (Cell Identity Factor, or CIF) for each cell by recursively evolving the parent cell’s CIF with a fate-dependent drift and Brownian noise:

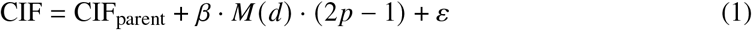

where CIF_parent_ is the latent state of the parent, *ε* ∼ 𝒩 (0, *σ*^2^)represents Brownian noise, and *β* controls the strength of the fate signal. The term (2*p* − 1) acts as a directional force: progenitors with a majority of Fate 1 descendants (*p* > 0.5) drift in the positive direction, while those favoring Fate 0 drift negatively. The function *M* (*d*) modulates this signal strength over time, modeling the biological reality that fate commitment becomes stronger as differentiation progresses (defined relative to commitment depth *D*):

- *M* (*d*) = 0.4 for *d* ≤ ⌊3*D*/7⌋ (early progenitors, weak signal)
- *M* (*d*) = 0.8 for ⌊3*D*/7⌋ < *d* ≤ *D* (intermediate commitment)
- *M* (*d*) = 1.2 for *d* > *D* (committed cells, strong signal)

### Phase 4: Gene expression readout

Each gene has associated sparse weight vectors that map CIFs to kinetic parameters. Using these kinetic parameters, we generate expression counts and add technical noise following the two-state kinetic model used by TedSim [Pan et al., 2022].

## 3 Experiments

We evaluate whether LineageSim’s fate bias is detectable: if progenitor expression encodes fate information, a classifier trained on it should predict terminal fate above chance level.

Using LineageSim, we generated a lineage tree with 8,192 cells and 1,000 genes. We sampled progenitors before commitment, assigned them binary labels based on fate majority, and balanced the classes by undersampling. We then trained a logistic regression classifier to predict fate from progenitor gene expression (Figure 1). Without fate bias (*β* = 0), we achieve 55.1% balanced accuracy, which is around chance level. This is expected when the expression carries no fate information. With fate bias enabled (*β* = 1), we achieve 68.3% balanced accuracy, well above chance level. This result confirms that the generated expression encodes meaningful fate information without making the prediction task trivial, mimicking the signal-to-noise ratio of real biological data. We further visualize the progenitor cells using UMAP (Figure 1b): with fate bias enabled, cells with similar fate probability *p* cluster together, confirming that the fate signal is encoded in the expression space.

**Figure 1.**
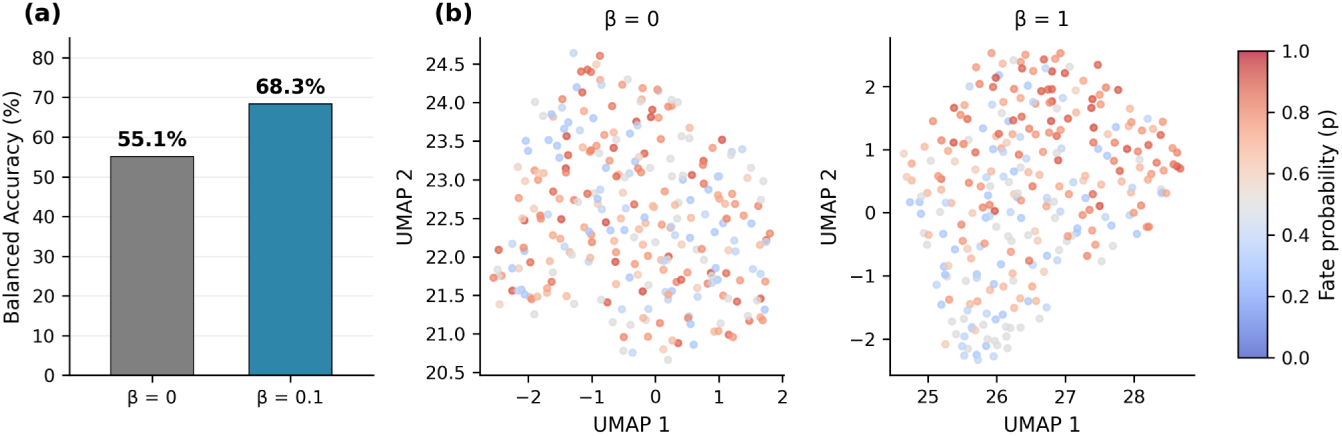
(a) Balanced accuracy of a logistic regression classifier that predicts binary fate from progenitor expression. Without fate bias (*β* = 0), balanced accuracy is 55.1%. With fate bias (*β* = 0.1), balanced accuracy reaches 68.3%, confirming detectable fate signals. (b) UMAP embedding of progenitor cells colored by fate probability *p*. When fate bias is present (*β* = 1), cells with similar fate probability *p* cluster together, demonstrating that fate bias is encoded in gene expression.

## 4 Conclusion

We introduced LineageSim to bridge the gap between static simulations and the dynamic reality of cell differentiation. By formally modeling fate bias as a force in the latent space, LineageSim provides the ground truth necessary for validating predictive lineage algorithms, a capability missing from previous simulators. While this study validates the framework on binary differentiation, future work will extend this approach to complex multi-fate topologies and non-linear trajectories. We release LineageSim as an open-source package, providing the community with a tool to rigorously benchmark the next generation of fate prediction models.

## References

Lennart Kester and Alexander van Oudenaarden. Single-cell transcriptomics meets lineage tracing. Cell Stem Cell, 23(2):166–179, 2018. doi: 10.1016/j.stem.2018.04.014.

Xinhai Pan, Hechen Li, and Xiuwei Zhang. Tedsim: temporal dynamics simulation of single-cell rna sequencing data and cell division history. Nucleic Acids Research, 50(8):4272–4288, 2022. doi: 10.1093/nar/gkac235.

Mehrshad Sadria and Thomas M. Bury. Fatenet: an integration of dynamical systems and deep learning for cell fate prediction. Bioinformatics, 40(9):btae525, 2024. doi: 10.1093/bioinformatics/btae525.

Mehrshad Sadria and Vasu Swaroop. Discovering governing equations of biological systems through representation learning and sparse model discovery. NAR Genomics and Bioinformatics, 7(2):qaf048, 2025. doi: 10.1093/nargab/lqaf048.

Mehrshad Sadria, Anita Layton, Sidhartha Goyal, and Gary D. Bader. Fatecode enables cell fate regulator prediction using classification-supervised autoencoder perturbation. Cell Reports Methods, 4(7):100819, 2024. doi: 10.1016/j.crmeth.2024.100819.

Shou-Wen Wang, Michael J. Herriges, Kilian Hurley, Darrell N. Kotton, and Allon M. Klein. Cospar identifies early cell fate biases from single-cell transcriptomic and lineage information. Nature Biotechnology, 40(7):1066–1074, 2022. doi: 10.1038/s41587-022-01209-1.

